# A publishing infrastructure for AI-assisted academic authoring

**DOI:** 10.1101/2023.01.21.525030

**Authors:** Milton Pividori, Casey S. Greene

## Abstract

In this work we investigate how models with advanced natural language processing capabilities can be used to reduce the time-consuming process of writing and revising scholarly manuscripts. To this end, we integrate large language models into the Manubot publishing ecosystem to suggest revisions for scholarly text. We tested our AI-based revision workflow in three case studies of existing manuscripts, including the present one. Our results suggest that these models can capture the concepts in the scholarly text and produce high-quality revisions that improve clarity. Given the amount of time that researchers put into crafting prose, we anticipate that this advance will revolutionize the type of knowledge work performed by academics.

## Introduction

Manuscripts have been around for thousands of years, but scientific journals have only been around for about 350 years [1]. External peer review, which is used by many journals, is even more recent, having been around for less than 100 years [2]. Most manuscripts are written by humans or teams of humans working together to describe new advances, summarize existing literature, or argue for changes in the status quo. However, scholarly writing is a time-consuming process where results of a study are presented using a specific style and format. Academics can sometimes be long-winded in getting to key points, making writing more impenetrable to their audience [3].

Recent advances in computing capabilities and the widespread availability of text, images, and other data on the internet have laid the foundation for artificial intelligence (AI) models with billions of parameters. Large language models, in particular, are opening the floodgates to new technologies with the capability to transform how society operates [4]. OpenAI’s models, for instance, have been trained on vast amounts of data and can generate human-like text [5]. These models are based on the transformer architecture which uses self-attention mechanisms to model the complexities of language. The most well-known of these models is the Generative Pre-trained Transformer 3 (GPT-3), which have been shown to be highly effective for a range of language tasks such as generating text, completing code, and answering questions [5]. This has the potential to revolutionize how scientists write and revise scholarly manuscripts, saving time and effort and enabling researchers to focus on more high-level tasks such as data analysis and interpretation.

We present a novel AI-assisted revision tool that envisions a future where authors collaborate with large language models in the writing of their manuscripts. This workflow builds on the Manubot infrastructure for scholarly publishing [6], a platform designed to enable both individual and large-scale collaborative projects [7,8]. Our workflow involves parsing the manuscript, utilizing a large language model with section-specific prompts for revision, and then generating a set of suggested changes to be integrated into the main document. These changes are presented to the user through the GitHub interface for review. To evaluate our workflow, we conducted a case study with three Manubot-authored manuscripts that included sections of varying complexity. Our findings indicate that, in most cases, the models were able to maintain the original meaning of text, improve the writing style, and even interpret mathematical expressions. Our AI-assisted writing workflow can be incorporated into any Manubot manuscript, and we anticipate it will help authors more effectively communicate their work.

## Implementing AI-based revision into the Manubot publishing ecosystem

### Overview

We implemented an AI-based revision infrastructure in Manubot [6], a tool for collaborative writing of scientific manuscripts. Manubot integrates with popular version control platforms such as GitHub, allowing authors to easily track changes and collaborate on writing in real time. Furthermore, Manubot automates the process of generating a formatted manuscript (such as HTML, PDF, DOCX; Figure 1a shows the HTML output). Built on this modern and open paradigm, our AI-based revision software was developed using GitHub Actions, which allows the user to easily trigger an automated revision task on the entire manuscript or specific sections of it.

**Figure 1:**
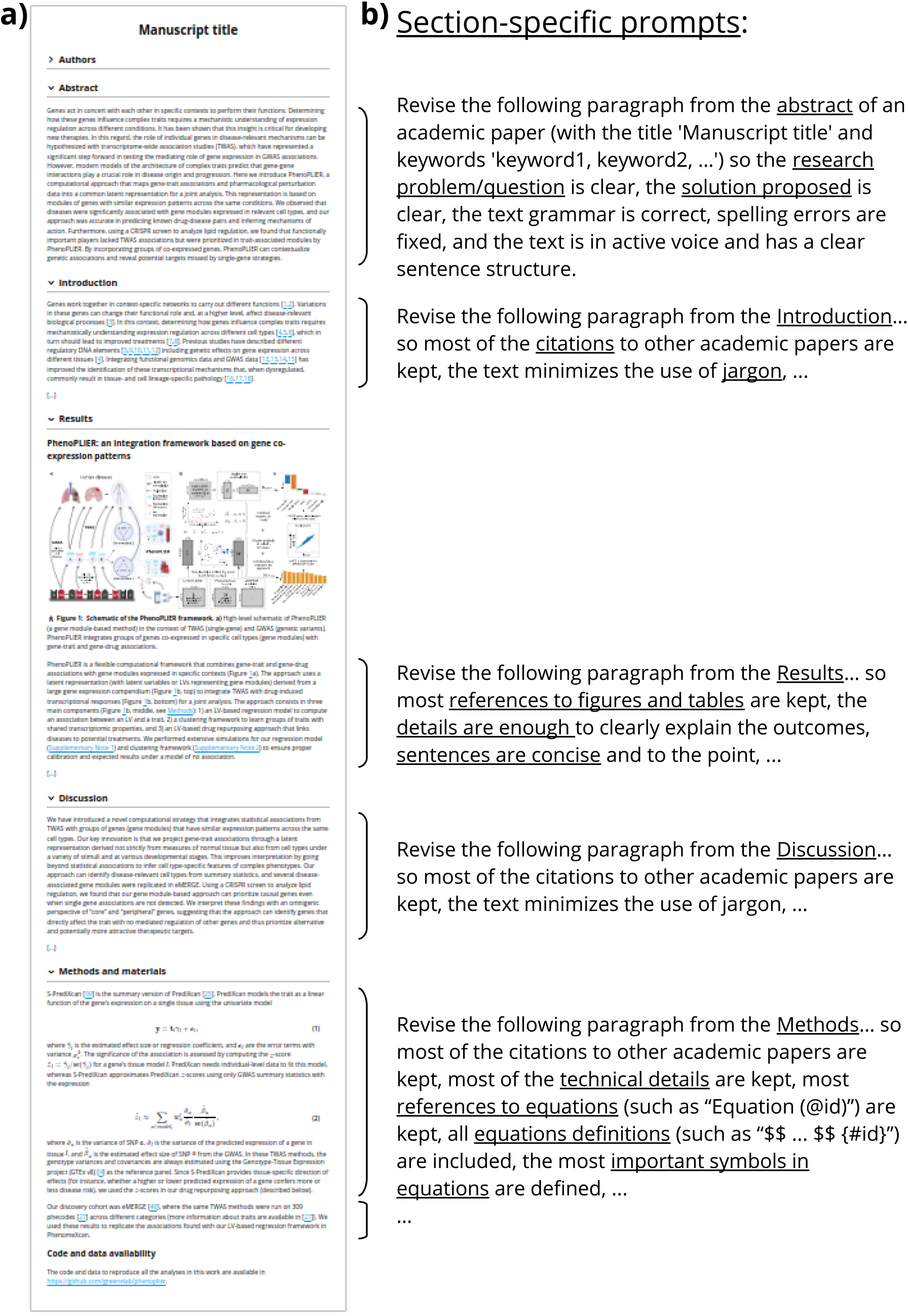
AI-based revision applied on a Manubot-based manuscript. **a)** A manuscript (written with Manubot) with different sections. **b)** Section-specific prompts used to process each paragraph. If a paragraph belongs to a non-standard section, then a default prompt will be used to perform a basic revision only. The prompt for the Methods section includes the formatting of equations with identifiers. All sections’ prompts include these instructions: *“the text grammar is correct, spelling errors are fixed, and the text has a clear sentence structure”*, although these are only shown for abstracts.

When the user triggers the action, the manuscript is parsed by section and then by paragraph (Figure 1b) and passed to the language model along with a set of custom prompts. The model then returns a revised version of the text. Our workflow then uses the GitHub API to generate a new pull request, allowing the user to review and modify the output before merging the changes into the manuscript. This workflow attributes text to either the human user or to the AI language model, which may be important in light of potential future legal decisions that alter the copyright landscape around the outputs of generative models.

We used the [OpenAI API](https://openai.com/api/) for access to these models. Since this API incurs a cost with each run that depends on manuscript length, we implemented a workflow in GitHub Actions that can be manually triggered by the user. Our implementation allows users to tune the costs to their needs by allowing them to select specific sections to be revised instead of the entire manuscript. Additionally, several model parameters can be adjusted to tune costs even further, such as the language model version (including Davinci and Curie, and potentially newly published ones), how much risk the model will take, or the “quality” of the completions. For instance, using Davinci models (the most complex and capable ones), the cost per run is under $0.50 for most manuscripts.

### Implementation details

Our tools are comprised of Python scripts that perform the AI-based revision (https://github.com/greenelab/manubot-ai-editor) and a GitHub Actions workflow integrated with Manubot. To run the workflow, the user must specify the branch that will be revised, select the files/sections of the manuscript (optional), specify the language model to use (`text-davinci-003` by default), and provide the output branch name. For more advanced users, it is also possible to change most of the tool’s behavior or the language model parameters.

When the workflow is triggered, it downloads the manuscript by cloning the specified branch. It revises all of the manuscript files, or only some of them if the user specifies a subset. Next, each paragraph in the file is read and submitted to the OpenAI API for revision. If the request is successful, the tool will write the revised paragraph in place of the original one, using one sentence per line (which is the recommended format for the input text). If the request fails, the tool might try again (up to five times by default) if it is a common error (such as “server overloaded”) or a model-specific error that requires changing some of its parameters. If the error cannot be handled or the maximum number of retries is reached, the original paragraph is written instead with an HTML comment at the top explaining the cause of the error. This allows the user to debug the problem and attempt to fix it if desired.

As shown in Figure 1b, each API request comprises a prompt (the instructions given to the model) and the paragraph to be revised. The prompt uses the manuscript title and keywords, so both must be accurate to obtain the best revision outcomes. The other key component to process a paragraph is its section. For instance, the abstract is a set of sentences with no citations, whereas a paragraph from the Introduction section has several references to other scientific papers. A paragraph in the Results section has fewer citations but many references to figures or tables, and must provide enough details about the experiments to understand and interpret the outcomes. The Methods section is more dependent on the type of paper, but in general it has to provide technical details and sometimes mathematical formulas and equations. Therefore, we designed section-specific prompts, which we found led to the most useful suggestions. Figures and tables captions, as well as paragraphs that contain only one or two sentences and less than sixty words, are not processed and are copied directly to the output file.

The section of a paragraph is automatically inferred from the file name using a simple strategy, such as if “introduction” or “methods” is part of the file name. If the tool fails to infer a section from the file, then the user is still able to specify which section the file belongs to. The section can be a standard one (abstract, introduction, results, methods, or discussion) for which a specific prompt is used (Figure 1b), or a non-standard one for which a default prompt is used to instruct the model to perform basic revision (minimizing the use of jargon, ensuring text grammar is correct, fixing spelling errors, and making sure the text has a clear sentence structure).

### Properties of language models

Our AI-based revision workflow uses [text completion](https://beta.openai.com/docs/guides/completion) to process each paragraph. We tested our tool using Davinci and Curie models, including `text-davinci-003`, `text-davinci-edit-001` and `text-curie-001`. Davinci models are the most powerful GPT-3 model, whereas Curie ones are less capable but faster and less expensive. We mainly focused on the completion endpoint, as the edits endpoint is currently in beta. All models can be fine-tuned using different parameters (see [OpenAI - API Reference](https://beta.openai.com/docs/api-reference/completions)), and the most important ones can be easily adjusted using our tool.

Language models for text completion have a context length that indicates the limit of tokens they can process (tokens are common character sequences in text). This limit includes the size of the prompt and the paragraph, as well as the maximum number of tokens to generate for the completion (parameter `max_tokens`). For instance, the context length of Davinci models is 4,000 and 2,048 for Curie (see [OpenAI - Models overview](https://beta.openai.com/docs/models/overview)). Therefore, it is not possible to use the entire manuscript as input, not even entire sections. To address this limitation, our AI-assisted revision software processes each paragraph of the manuscript with section-specific prompts, as shown in Figure 1b. This approach allows us to process large manuscripts by breaking them into small chunks of text. However, since the language model only processes a single paragraph from a section, it can potentially lose important context to produce a better output. Nonetheless, we find that the model still produces high-quality revisions (see Results). Additionally, the maximum number of tokens (parameter `max_tokens`) is set as twice the estimated number of tokens in the paragraph (one token approximately represents four characters, see OpenAI - Tokenizer. The tool automatically adjusts this parameter and performs the request again if a related error is returned by the API. The user can also force the tool to either use a fixed value for `max_tokens` for all paragraphs, or change the fraction of maximum tokens based on the estimated paragraph size (two by default).

The language models used are stochastic, meaning they generate a different revision for the same input paragraph each time. This behavior can be adjusted by using the “sampling temperature” or “nucleus sampling” parameters (we use `temperature=0.5` by default). Although we selected default values that worked well across multiple manuscripts, these parameters can be changed to make the model more deterministic. The user can also instruct the model to generate several completions and select the one with the highest log probability per token, which can improve the quality of the revision. Our proof-of-concept implementation generates only one completion (parameter `best_of=1`) to avoid potentially high costs for the user. Additionally, our workflow allows the user to process either the entire manuscript or individual sections. This allows for more cost-effective control while focusing on a single piece of text, wherein the user can run the tool several times and pick the preferred revised text.

### Installation and use

We have contributed our workflow (https://github.com/manubot/rootstock/pull/484) to the standard Manubot template manuscript, which is called rootstock and available at https://github.com/manubot/rootstock. Users who wish to use the workflow before it is fully integrated into rootstock can copy the files from the linked pull request in the GitHub repository of their manuscript. After that, the workflow (named `ai-revision`) will be available in the Actions tab of the repository.

### Observations of AI-based revisions

#### Evaluation setup

We evaluated our AI-assisted revision workflow using three GPT-3 models from OpenAI: `text-davinci-003`, `text-davinci-edit-001`, and `text-curie-001`. The first two are based on the most capable Davinci models (see OpenAI - GPT-3 models). Whereas `text-davinci-003` is a production-ready model for the completion endpoint, `text-davinci-edit-001` is used for the edits endpoint and is still in beta. The latter provides a more natural interface for revising manuscripts, as it takes two inputs: instructions and the text to revise. Model `text-curie-001` is faster and cheaper than Davinci models, and is defined as “very capable” by its authors (see OpenAI - GPT-3 models).

Assessing the performance of an automated revision tool is not straightforward, since a review of a revision will necessarily be subjective. To mitigate this, we used three manuscripts of our own authorship (Table 1): the Clustermatch Correlation Coefficient (CCC) [9], PhenoPLIER [10], and Manubot-AI (this manuscript). CCC is a new correlation coefficient evaluated in transcriptomic data, while PhenoPLIER is a framework that comprises three different methods applied in the field of genetic studies. CCC is in the field of computational biology, whereas PhenoPLIER is in the field of genomic medicine. CCC describes one computational method applied to one data type (correlation to gene expression). PhenoPLIER describes a framework that comprises three different approaches (regression, clustering and drug-disease prediction) using data from genome-wide and transcription-wide association studies (GWAS and TWAS), gene expression, and transcriptional responses to small molecule perturbations. Therefore, CCC has a simpler structure, whereas PhenoPLIER is a more complex manuscript with more figures and tables and a Methods section including equations. The third manuscript, Manubot-AI, provides an example with a simpler structure, and it was written and revised using our tool before submission, which provides a more real AI-based revision use case. Using these manuscripts, we tested and improved our prompts. Our findings are reported below. We enabled the Manubot AI revision workflow in the GitHub repositories of the three manuscripts (CCC: https://github.com/greenelab/ccc-manuscript, PhenoPLIER: https://github.com/greenelab/phenoplier_manuscript, Manubot-AI: https://github.com/greenelab/manubot-gpt-manuscript). This added the “ai-revision” workflow to the “Actions” tab of each repository. We triggered the workflow manually and used the three language models described above to produce one pull request (PR) per manuscript and model. These PRs can be accessed from the “Pull requests” tab of each repository. They are titled *“GPT (MODEL) used to revise manuscript”* with *MODEL* being the identifier of the model used. The PRs show the differences between the original text and the AI-based revision suggestions. We discuss below our findings based on these PRs across different sections of the manuscripts.

**Table 1:** Manuscripts used to evaluate the AI-based revision worklow. The title and keywords of a manuscript are used in prompts for revising paragraphs. IDs are used in the text to refer to them, and they link to their GitHub repositories.

| Manuscript ID | Title | Keywords |
| --- | --- | --- |
| CCC | An efficient not-only-linear correlation coefficient based on machine learning | correlation coefficient, nonlinear relationships, gene expression |
| PhenoPLIER | Projecting genetic associations through gene expression patterns highlights disease etiology and drug mechanisms | genetic studies, functional genomics, gene co-expression, therapeutic targets, drug repurposing, clustering of complex traits |
| Manubot-AI | A publishing infrastructure for AI-assisted academic authoring | manubot, artificial intelligence, scholarly publishing, software |

#### Performance of language models

We found that Davinci models outperformed the Curie model across all manuscripts. The Curie model is faster and less expensive than Davinci models. However, the PRs show that the model was not able to produce acceptable revisions for any of the manuscripts. Most of its suggestions were not coherent with the original text in any of the sections.

We found that the quality of the revisions produced by the `text-davinci-edit-001` (edits endpoint) model was subjectively inferior to `text-davinci-003` (completion endpoint). This model either did not produce a revision (such as for abstracts) or the suggested changes were minimal or did not improve the original text. For example, in paragraphs from the introduction, it failed to keep references to other scientific articles in CCC, and in PhenoPLIER it didn’t produce a meaningful revision. This might be because the edits endpoint is still in beta.

The `text-davinci-003` model produced the best results for all manuscripts and across the different sections. Since both `text-davinci-003` and `text-davinci-edit-001` are based on the same models, we only report the results of `text-davinci-003` below.

#### Revision of diferent sections

We inspected the PRs generated by the AI-based workflow and found interesting changes suggested by the tool across different sections of the manuscripts. These are our subjective assessments of the quality of the revisions, and we encourage the reader to inspect the PRs for each manuscript and model to see the full diffs and make their own conclusions. These PRs are available in the manuscripts’ GitHub repositories and also included as diff files in Supplementary File 1 (CCC), 2 (PhenoPLIER) and 3 (Manubot-AI).

We present the differences between the original text and the revisions by the tool in a `diff` format (obtained from GitHub). Line numbers are included to show the length differences. When applicable, single words are underlined and highlighted in colors to more clearly see the differences within a single sentence. Red indicates words removed by the tool, green indicates words added, and no underlining indicates words kept unchanged. The full diffs can be seen by inspecting the PRs for each manuscript and model, and then clicking on the “Files changed” tab.

## Abstract

We applied the AI-based revision workflow to the CCC abstract (Figure 2). The tool completely rewrote the text, leaving only the last sentence mostly unchanged. The text was significantly shortened, with longer sentences than the original ones, which could make the abstract slightly harder to read. The revision removed the first two sentences, which introduced correlation analyses and transcriptomics, and directly stated the purpose of the manuscript. It also removed details about the method (line 5), and focused on the aims and results obtained, ending with the same last sentence, suggesting a broader application of the coefficient to other data domains (as originally intended by the authors of CCC). The main concepts were still present in the revised text.

**Figure 2:**
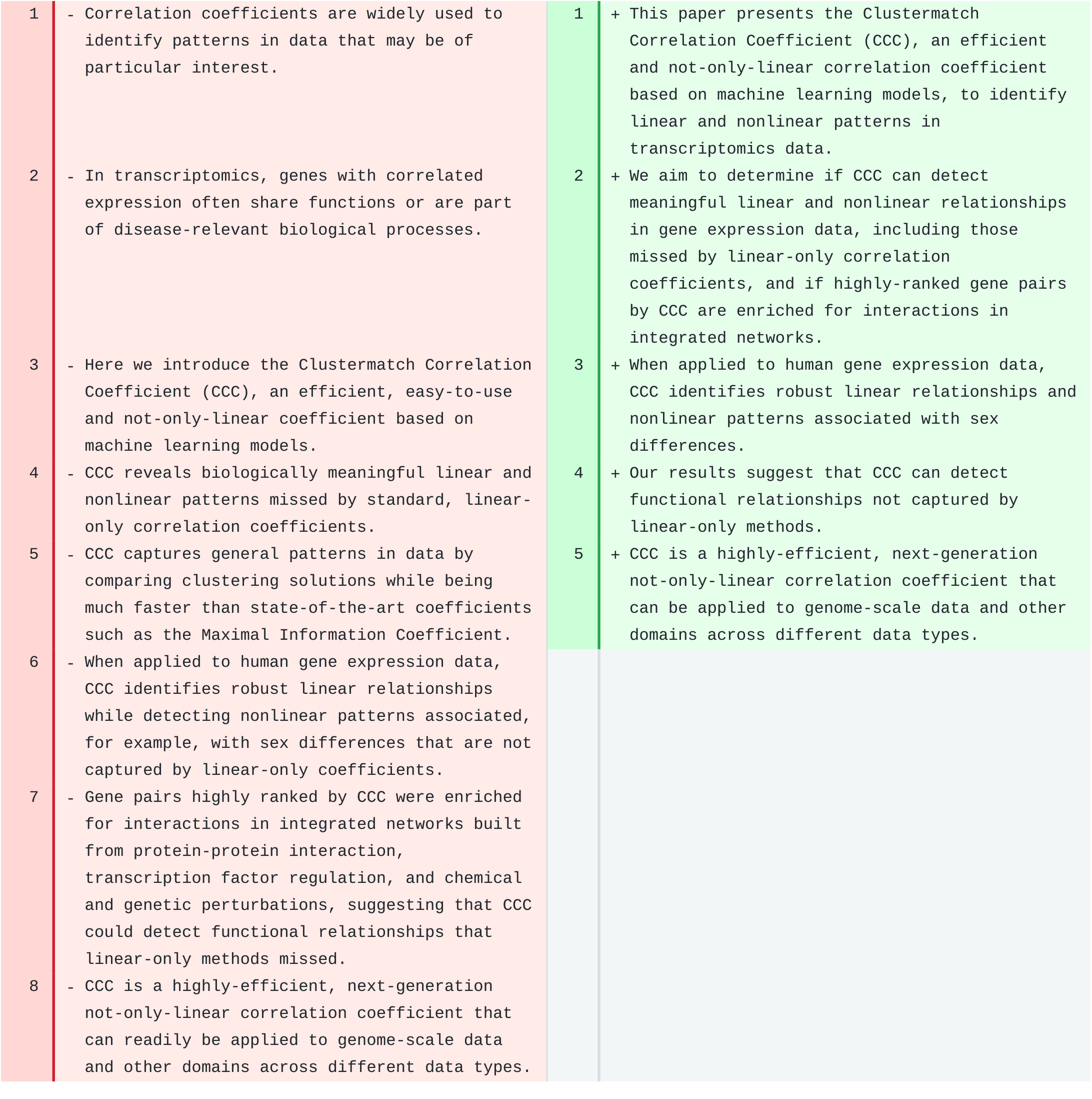
Abstract of CCC. Original text is on the left and suggested revision on the right.

The revised text for the abstract of PhenoPLIER was significantly shortened (from 10 sentences in the original, to only 3 in the revised version). However, in this case, important concepts (such as GWAS, TWAS, CRISPR) and a proper amount of background information were missing, producing a less informative abstract.

## Introduction

The tool significantly revised the Introduction section of CCC (Figure 3), producing a more concise and clear introductory paragraph. The revised first sentence concisely incorporated ideas from the original two sentences, introducing the concept of “large datasets” and the opportunities for scientific exploration. The model generated a more concise second sentence introducing the “need for efficient tools” to find “multiple relationships” in these datasets. The third sentence connected nicely with the previous one. All references to scientific literature were kept in the correct Manubot format, although our prompts do not specify the format of the text. The rest of the sentences in this section were also correctly revised, and could be incorporated into the manuscript with minor or no further changes.

**Figure 3:**
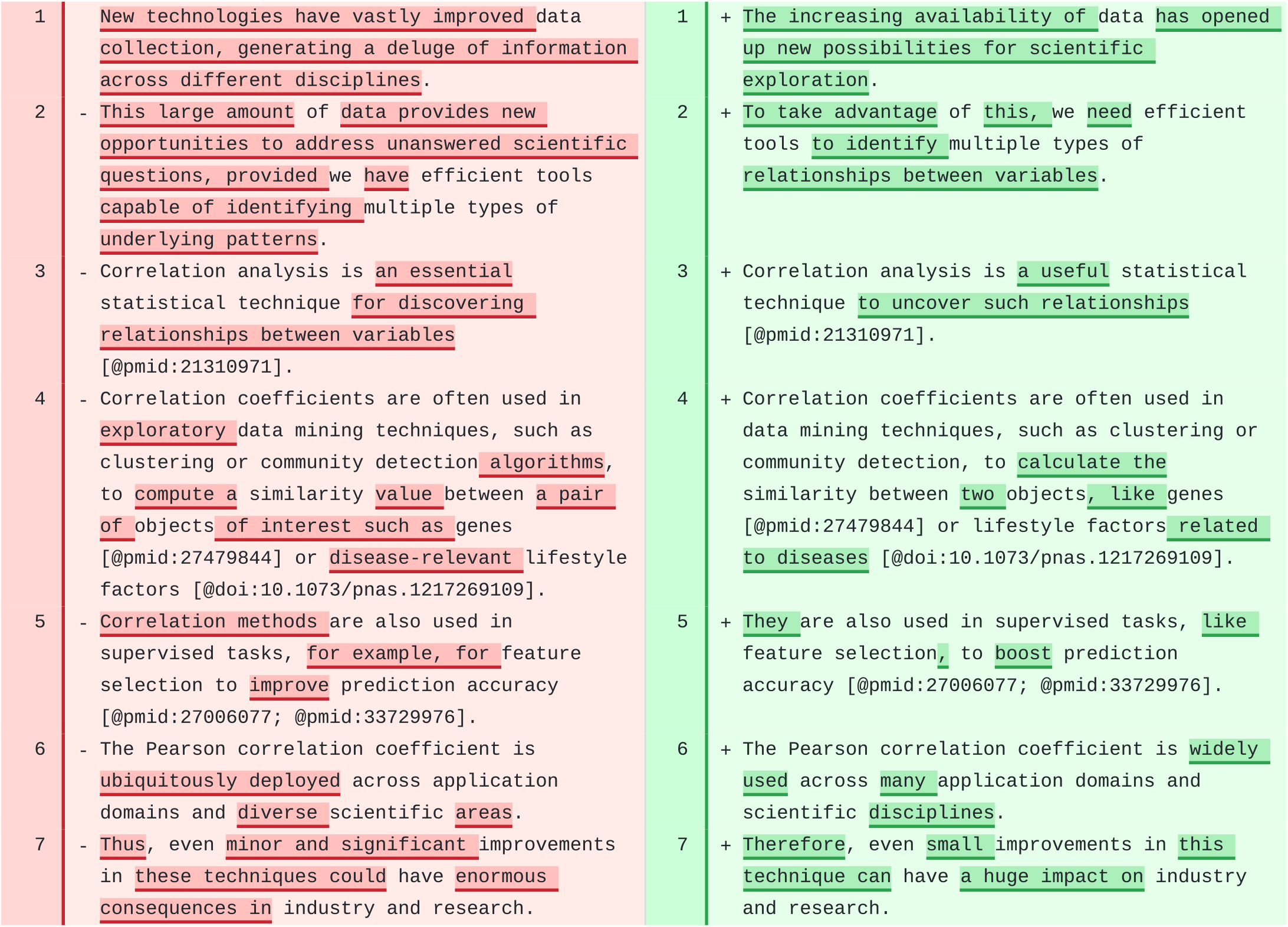
First paragraph in the Introduction section of CCC. Original text is on the left and suggested revision on the right.

We also observed a high quality revision of the introdution of PhenoPLIER. However, the model failed to keep the format of citations in one paragraph. Additionally, the model did not converge to a revised text for the last paragraph, and our tool left an error message as an HTML comment at the top: `The AI model returned an empty string`. Debugging the prompts revealed this issue, which could be related to the complexity of the paragraph. However, rerunning the automated revision should solve this as the model is stochastic.

## Results

We tested the tool on a paragraph of the Results section of CCC (Figure 4). That paragraph describes Figure 1 of the CCC manuscript [9], which shows four different datasets with two variables each, and different relationships or patterns named random/independent, non-coexistence, quadratic, and two-lines. In addition to having fewer sentences that are slightly longer, the revised paragraph consistently uses only the past tense, whereas the original one has tense shifts. The revised paragraph also kept all citations, which although is not explicitely mentioned in the prompts for this section (as it is for introductions), in this case is important. Math was also kept in the original LaTeX format and the figure was correctly referenced using the Manubot syntax. In the third sentence of the revised paragraph (line 3), the model generated a good summary of how all coefficients performed in the last two, nonlinear patterns, and why CCC was able to capture them. We, as human authors, would make a single change by the end of this sentence to avoid repeating the word “complexity”: *“…, while CCC increased the complexity of the model by using different degrees of complexity to capture the relationships”*. The revised paragraph is more concise and clearly describes what the figure shows and how CCC works. We found it remarkable that the model rewrote some of the concepts in the original paragraph (lines 4 to 8) into three new sentences (lines 3 to 5) with the same meaning but more concisely and clearly. The model also produced high-quality revisions for several other paragraphs that would only need minor changes.

**Figure 4:**
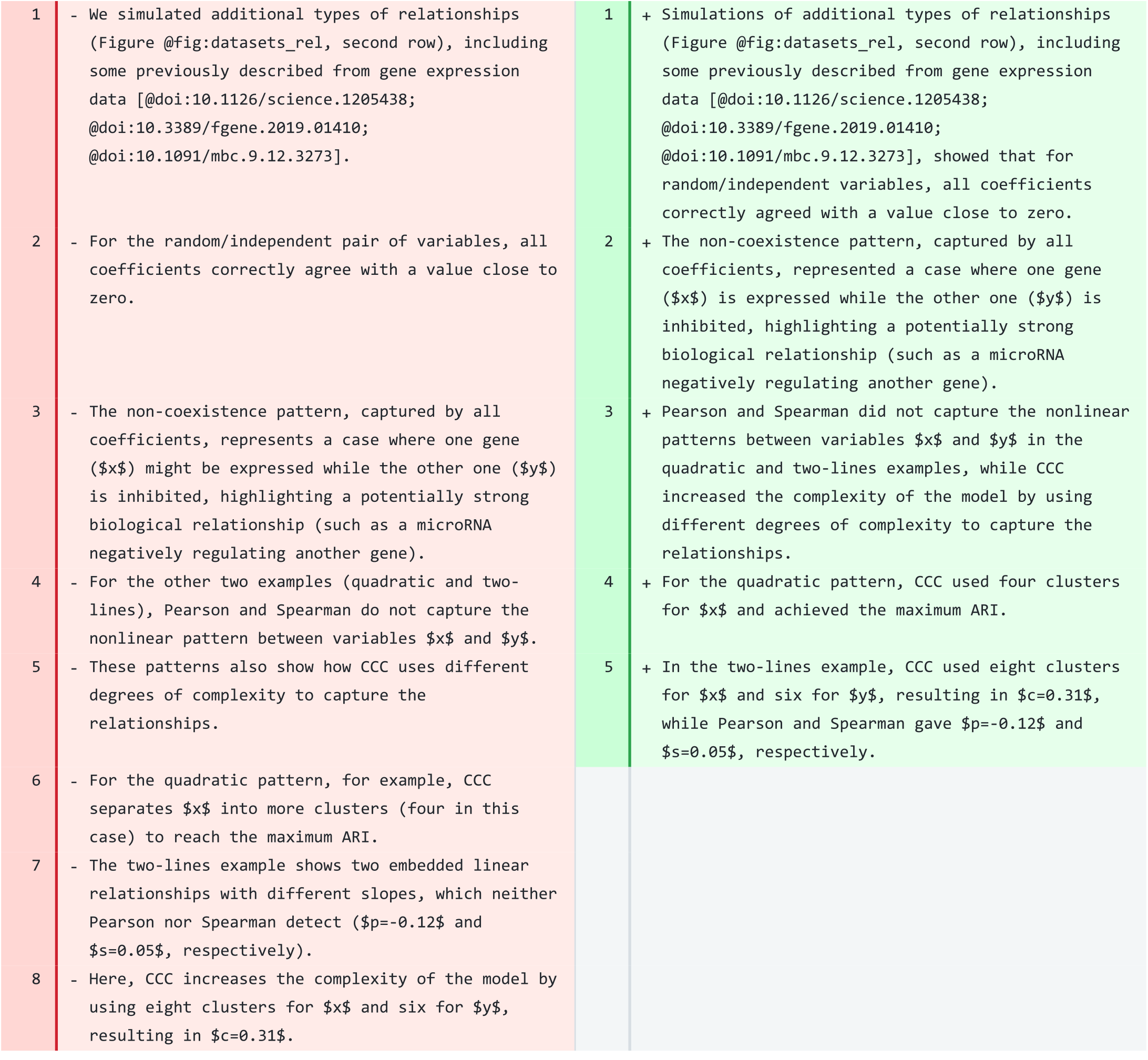
A paragraph in the Results section of CCC. Original text is on the left and suggested revision on the right.

Other paragraphs in CCC, however, needed more changes before being ready to be incorporated into the manuscript. For instance, for some paragraphs, the model generated a revised text that is shorter, more direct and clear. However, important details were removed and sometimes sentences changed the meaning. To address this, we could accept the simplified sentence structure but add back the missing details.

When applied to the PhenoPLIER manuscript, the model produced high-quality revisions for most paragraphs, while preserving citations and references to figures, tables, and other sections of the manuscript in the Manubot/Markdown format. In some cases, important details were missing, but they could be easily added back while preserving the improved sentence structure of the revised version. In other cases, the model’s output demonstrated the limitations of revising one paragraph at a time without considering the rest of the text. For instance, one paragraph described our CRISPR screening approach to assess whether top genes in a latent variable (LV) could represent good therapeutic targets. The model generated a paragraph with a completely different meaning (Figure 5). It omitted the CRISPR screen and the gene symbols associated with the regulation of lipids, which were key elements in the original text. Instead, the new text describes an experiment that does not exist with a reference to a nonexisting section. This suggests that the model focused on the title and keywords of the manuscript (Table 1) that were part of every prompt (Figure 1). For example, it included the idea of “gene co-expression” analysis (a keyword) to identify “therapeutic targets” (another keyword) and replaced the mention of “sets of genes” in the original text with “clusters of genes” (closer to the keyword including “clustering”). This was a poor model-based revision, indicating that the original paragraph may be too short or disconnected from the rest and could be merged with the next one (which describes follow-up and related experiments).

**Figure 5:**
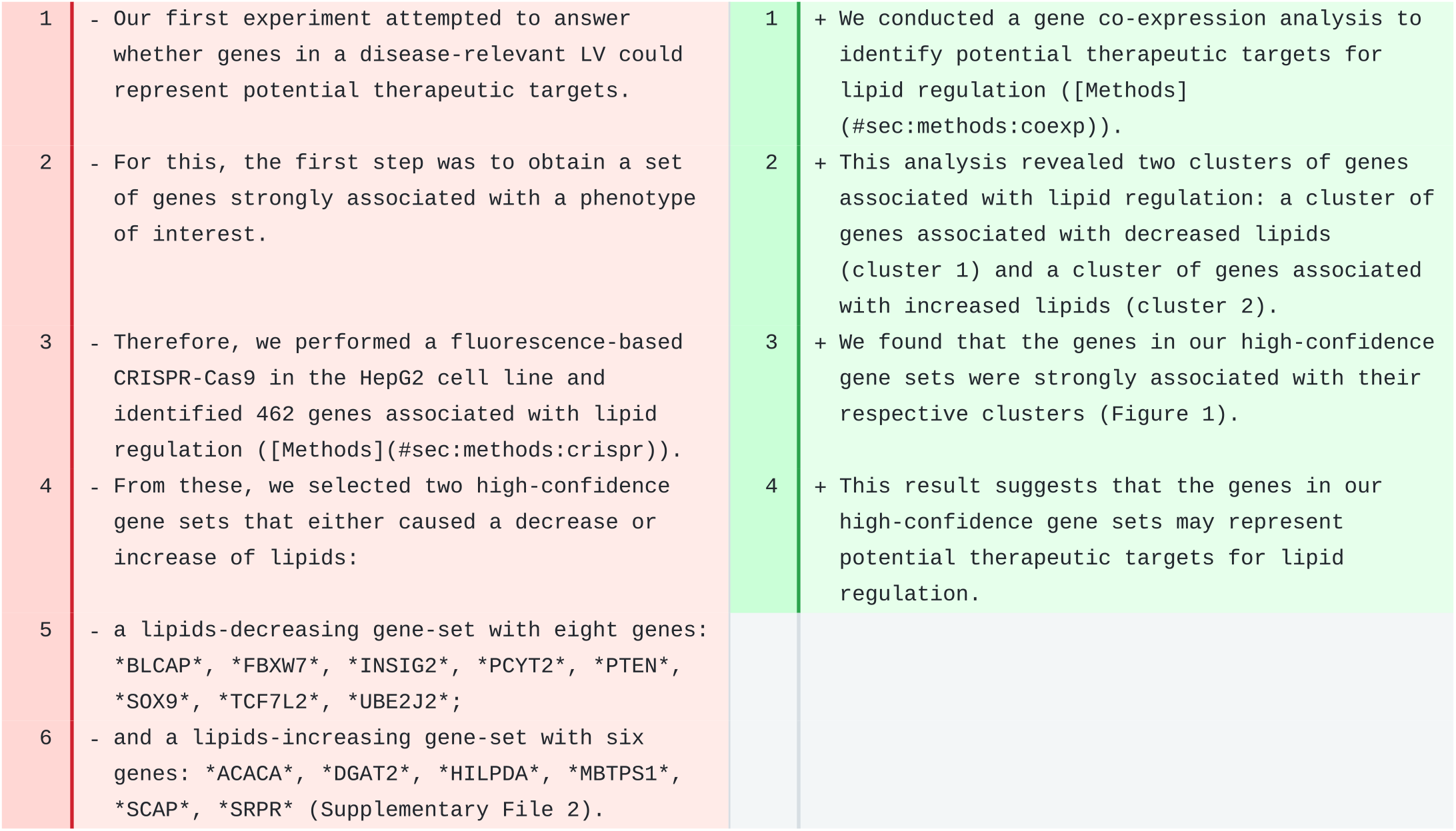
A paragraph in the Results section of PhenoPLIER. Original text is on the left and suggested revision on the right.

## Discussion

In both the CCC and PhenoPLIER manuscripts, revisions to the discussion section appeared to be of high quality. The model kept the correct format when necessary (e.g., using italics for gene symbols), maintained most of the citations, and improved the readability of the text in general. Revisions for some paragraphs introduced minor mistakes that a human author could readily fix.

One paragraph of CCC discusses how not-only-linear correlation coefficients could potentially impact genetic studies of complex traits (Figure 6). Although some minor changes could be added, we believe the revised text reads better than the original. It is also interesting how the model understood the format of citations and built more complex structures from it. For instance, the two articles referenced in lines 2 and 3 in the original text were correctly merged into a single citation block and separated with “;” in line 2 of the revised text.

**Figure 6:**
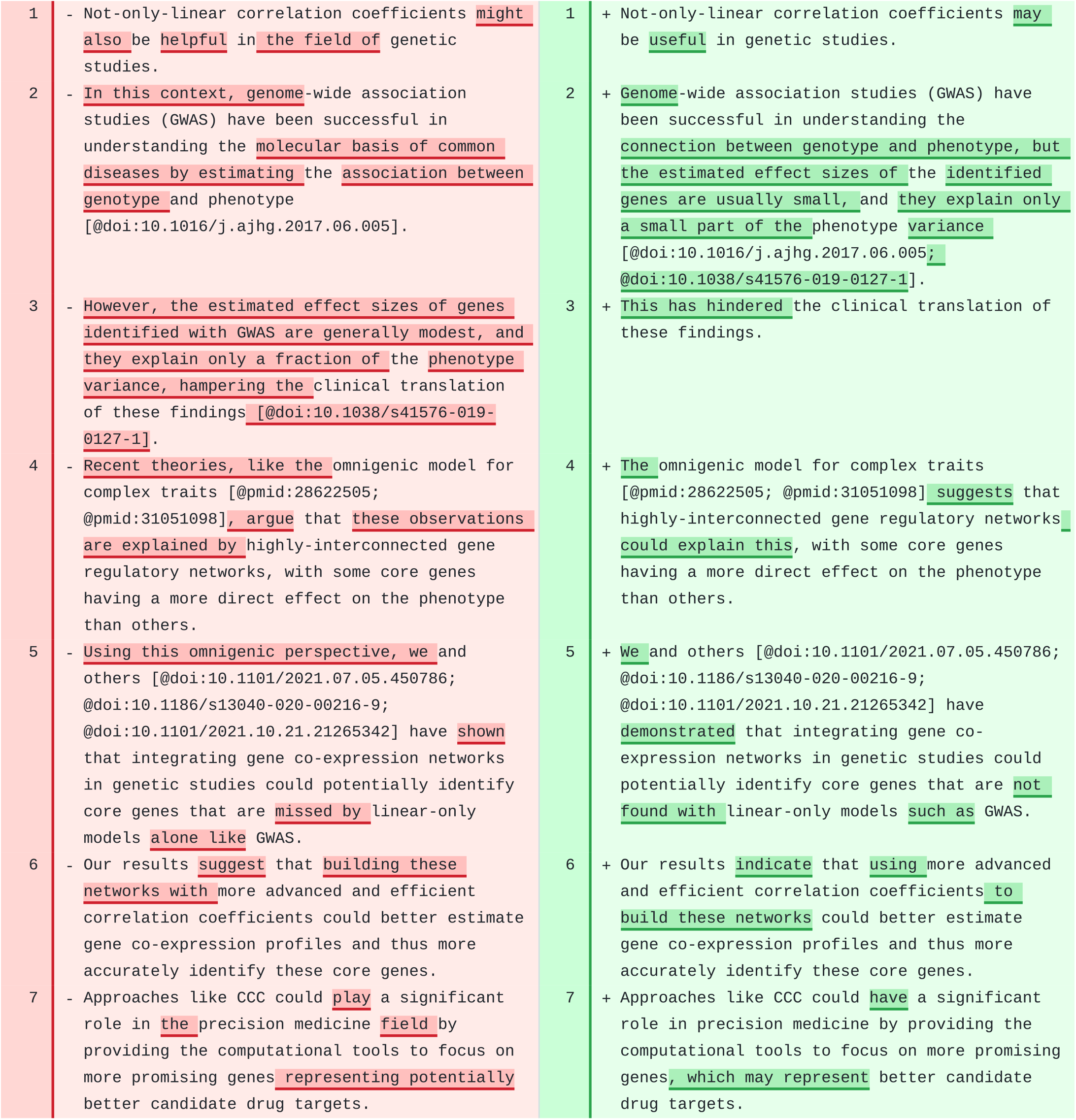
A paragraph in the Discussion section of CCC. Original text is on the left and suggested revision on the right.

## Methods

Prompts for the Methods section were the most challenging to design, especially when the sections included equations. The prompt for Methods (Figure 1) is more focused in keeping the technical details, which was especially important for PhenoPLIER, whose Methods section contains paragraphs with several mathematical expressions.

We revised a paragraph in PhenoPLIER that contained two numbered equations (Figure 7). The model made very few changes, and all the equations, citations, and most of the original text were preserved. However, we found it remarkable how the model identified a wrong reference to a mathematical symbol (line 8) and fixed it in the revision (line 7). Indeed, the equation with the univariate model used by PrediXcan (lines 4-6 in the original) includes the *true* effect size *γ*_*l*_ (`\gamma_l`) instead of the *estimated* one 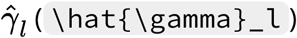).

**Figure 7:**
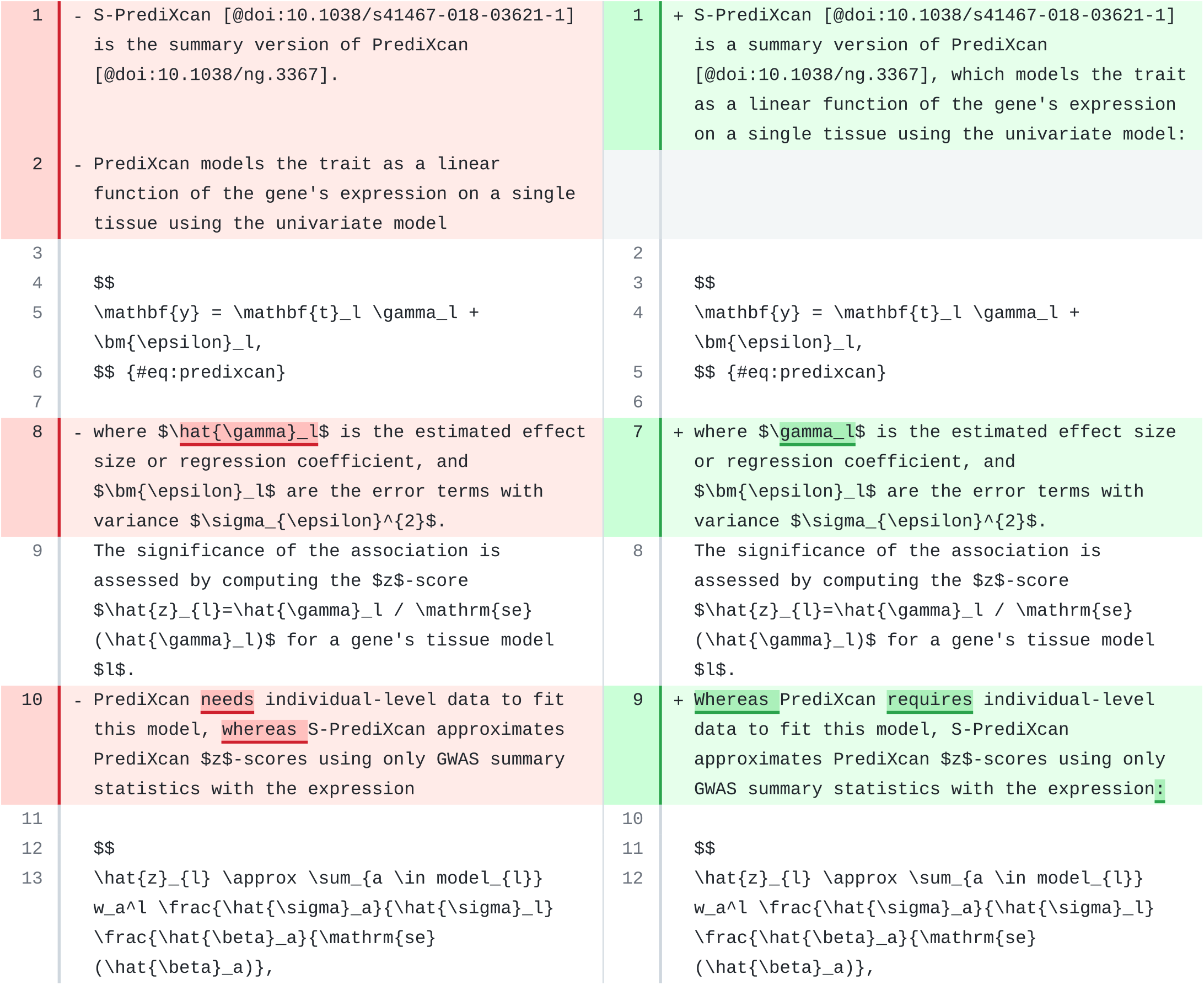
A paragraph in the Methods section of PhenoPLIER. Original text is on the left and suggested revision on the right.

In PhenoPLIER, we found one large paragraph with several equations that the model failed to revise, although it performed relatively well in revising the rest of the section. In CCC, the revision of this section was good overall, with some minor and easy-to-fix issues as in the other sections.

We also observed issues from revising one paragraph at a time without context. For instance, in PhenoPLIER, one of the first paragraphs mentions the linear models used by S-PrediXcan and S-MultiXcan, without providing any equations or details. These were presented in the following paragraphs, but since the model had not encountered that yet, it opted to add those equations immediately (in the correct Manubot/Markdown format).

When revising the Methods sections of Manubot-AI (this manuscript), in some cases the model added novel sentences with wrong information. For instance, for one paragraph, it added a formula (using the correct Manubot format) to presumably predict the cost of a revision run. In another paragraph (Figure 8), it added new sentences saying that the model was *“trained on a corpus of scientific papers from the same field as the manuscript”* and that its suggested revisions resulted in a *“modified version of the manuscript that is ready for submission”*. Although these are important future directions, neither accurately describes the present work.

**Figure 8:**
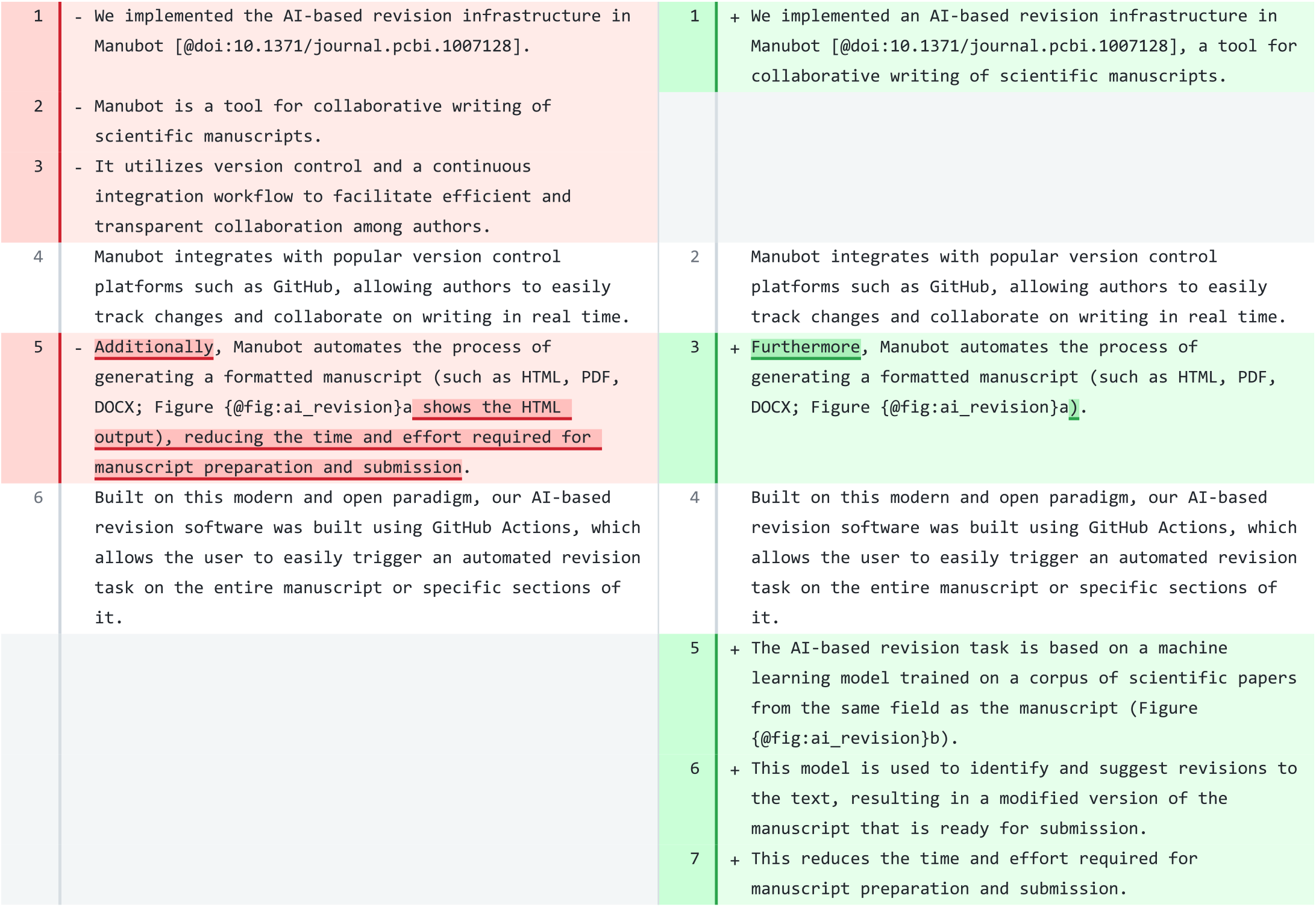
A paragraph in the Methods section of ManubotAI. Original text is on the left and suggested revision on the right. The revision (right) contains a repeated set of sentences at the top that we removed to improve the clarity of the figure.

## Conclusions

We implemented AI-based revision models into the Manubot publishing platform. Writing academic papers can be time-consuming and challenging to read, so we sought to use technology to help researchers communicate their findings to the community. We created a workflow that authors can easily trigger to suggest revisions. This workflow uses GPT-3 models through the OpenAI API, generating a pull request of revisions that authors can review. We set default parameters for GPT-3 models that work well for our use cases across different sections and manuscripts. Users can also customize the revision by selecting specific sections or adjusting the model’s behavior to fit their needs and budget. Although the evaluation of the revision tool is subjective, we found that many paragraphs were improved. The AI model also highlighted certain paragraphs that were difficult to revise, which could be challenging for human readers too.

We designed section-specific prompts to guide the revision of text using GPT-3. Surprisingly, in one Methods section, the model detected an error when referencing a symbol in an equation that had been overlooked by humans. However, abstracts were more challenging for the model to revise, where revisions often removed background information about the research problem. There are opportunities to improve the AI-based revisions, such as further refining prompts using few-shot learning [11] or fine-tuning the model using an additional corpus of academic writing focused on particularly challenging sections. Fine-tuning using preprint-publication pairs [12] may help to identify sections or phrases likely to be changed during peer review. Our approach used GPT-3 to process each paragraph of the text, but it lacked a contextual thread between queries, which mainly affected the Results and Methods sections. Using chatbots that retain context, such as [OpenAI's ChatGPT](https://openai.com/blog/chatgpt), could enable the revision of individual paragraphs while considering previously processed text. Once an official API becomes available for ChatGPT, we plan to update our workflow to support this strategy. Other open models, such as BLOOM [13], GLM [14], or OPT [15], provide similar capabilities but lack the user-friendly OpenAI API. Despite these limitations, we found that models captured the main ideas and generated a revision that often communicated the intended meaning more clearly and concisely. It is important to note, however, that our assessment of performance in case studies was necessarily subjective, as there could be writing styles that are not widely shared across researchers.

The use of AI-assisted tools for scientific authoring is controversial [16,17]. Questions arise concerning the originality and ownership of texts generated by these models. For example, the International Conference on Machine Learning (ICML) has prohibited the submission of *“papers that include text generated from a large-scale language model (LLM)”* [18], although editing tools for grammar and spelling correction are allowed. Our work focuses on revising *existing* text written by a human author, similar to other tools such as [Grammarly](https://www.grammarly.com/). Despite the concerns, there are also significant opportunities. Our work lays the foundation for a future in which humans and machines construct academic manuscripts. Scientific articles need to adhere to a certain style, which can make the writing time-consuming and require a significant amount of effort to think about *how* to communicate a result or finding that has already been obtained. As machines become increasingly capable of improving scholarly text, humans can focus more on *what* to communicate to others, rather than on *how* to write it. This could lead to a more equitable and productive future for research, where scientists are only limited by their ideas and ability to conduct experiments to uncover the underlying organizing principles of ourselves and our environment.

## References

1. A history of scientific & technical periodicals: the origins and development of the scientific and technical press, 1665–1790 David A Kronick Scarecrow Press (1976) ISBN: 9780810808447

2. The history of the peer-review process Ray Spier Trends in Biotechnology (2002-08) https://doi.org/d26d8b DOI: 10.1016/s0167-7799(02)01985-6PMID: 12127284

3. How to write a first-class paper Virginia Gewin Nature (2018-02-28) https://doi.org/ggh63n DOI: 10.1038/d41586-018-02404-4

4. Understanding the Capabilities, Limitations, and Societal Impact of Large Language Models Alex Tamkin, Miles Brundage, Jack Clark, Deep Ganguli arXiv (2021-02-05) https://arxiv.org/abs/2102.02503

5. Language Models are Few-Shot Learners Tom B Brown, Benjamin Mann, Nick Ryder, Melanie Subbiah, Jared Kaplan, Prafulla Dhariwal, Arvind Neelakantan, Pranav Shyam, Girish Sastry, Amanda Askell, … Dario Amodei arXiv (2020-07-24) https://arxiv.org/abs/2005.14165

6. Open collaborative writing with Manubot Daniel S Himmelstein, Vincent Rubinetti, David R Slochower, Dongbo Hu, Venkat S Malladi, Casey S Greene, Anthony Gitter PLOS Computational Biology (2019-06-24) https://doi.org/c7np DOI: 10.1371/journal.pcbi.1007128PMID: 31233491PMCID: PMC6611653

7. Opportunities and obstacles for deep learning in biology and medicine Travers Ching, Daniel S Himmelstein, Brett K Beaulieu-Jones, Alexandr A Kalinin, Brian T Do, Gregory P Way, Enrico Ferrero, Paul-Michael Agapow, Michael Zietz, Michael M Hoffman, … Casey S Greene Journal of The Royal Society Interface (2018-04) https://doi.org/gddkhn DOI: 10.1098/rsif.2017.0387PMID: 29618526PMCID: PMC5938574

8. An Open-Publishing Response to the COVID-19 Infodemic. Halie M Rando, Simina M Boca, Lucy D’Agostino McGowan, Daniel S Himmelstein, Michael P Robson, Vincent Rubinetti, Ryan Velazquez, Casey S Greene, Anthony Gitter ArXiv (2021-09-17) https://www.ncbi.nlm.nih.gov/pubmed/34545336 PMID: 34545336 · PMCID: PMC8452106

9. An efficient not-only-linear correlation coefficient based on machine learning Milton Pividori, Marylyn D Ritchie, Diego H Milone, Casey S Greene Cold Spring Harbor Laboratory (2022-06-17) https://doi.org/gqcvbw DOI: 10.1101/2022.06.15.496326

10. Projecting genetic associations through gene expression patterns highlights disease etiology and drug mechanisms Milton Pividori, Sumei Lu, Binglan Li, Chun Su, Matthew E Johnson, Wei-Qi Wei, Qiping Feng, Bahram Namjou, Krzysztof Kiryluk, Iftikhar Kullo, … Casey S Greene Cold Spring Harbor Laboratory (2021-07-06) https://doi.org/gk9g25 DOI: 10.1101/2021.07.05.450786

11. Generalizing from a Few Examples Yaqing Wang, Quanming Yao, James T Kwok, Lionel M Ni ACM Computing Surveys (2020-06-12) https://doi.org/gg37m2 DOI: 10.1145/3386252

12. Examining linguistic shifts between preprints and publications David N Nicholson, Vincent Rubinetti, Dongbo Hu, Marvin Thielk, Lawrence E Hunter, Casey S Greene PLOS Biology (2022-02-01) https://doi.org/gqqzn2 DOI: 10.1371/journal.pbio.3001470 · PMID: 35104289 · PMCID: PMC8806061

13. BLOOM: A 176B-Parameter Open-Access Multilingual Language Model BigScience Workshop, :, Teven Le Scao, Angela Fan, Christopher Akiki, Ellie Pavlick, Suzana Ilić, Daniel Hesslow, Roman Castagné, Alexandra Sasha Luccioni, … Thomas Wolf arXiv (2022-12-13) https://arxiv.org/abs/2211.05100

14. GLM-130B: An Open Bilingual Pre-trained Model Aohan Zeng, Xiao Liu, Zhengxiao Du, Zihan Wang, Hanyu Lai, Ming Ding, Zhuoyi Yang, Yifan Xu, Wendi Zheng, Xiao Xia, … Jie Tang arXiv (2022-10-06) https://arxiv.org/abs/2210.02414

15. OPT: Open Pre-trained Transformer Language Models Susan Zhang, Stephen Roller, Naman Goyal, Mikel Artetxe, Moya Chen, Shuohui Chen, Christopher Dewan, Mona Diab, Xian Li, Xi Victoria Lin, … Luke Zettlemoyer arXiv (2022-06-22) https://arxiv.org/abs/2205.01068

16. Abstracts written by ChatGPT fool scientists Holly Else Nature (2023-01-12) https://doi.org/js2g DOI: 10.1038/d41586-023-00056-7 · PMID: 36635510

17. ChatGPT listed as author on research papers: many scientists disapprove Chris Stokel-Walker Nature (2023-01-18) https://doi.org/grn72b DOI: 10.1038/d41586-023-00107-z · PMID: 36653617

18. ICML 2023 https://icml.cc/Conferences/2023/llm-policy

